# Assessment of antimicrobial activity of ethanolic and aqueous extracts of *Aesculus hippocastanum* L. (horse chestnut) bark against bacteria isolated from urine of patients diagnosed positive to urinary tract infections

**DOI:** 10.1101/2021.08.23.457193

**Authors:** K. Khar’kov Yuriy, Mbarga Manga Joseph Arsene, V. Martynenkova Aliya, V. Podoprigora Irina, G. Volina Elena

## Abstract

The search for new antimicrobials is essential to address the worldwide issue of antibiotic resistance which affects all areas requiring the use of antibiotics including the management of diseases such as urinary tract infections (UTIs).

**Aim:** To assess the antimicrobial activity of ethanolic and aqueous extract of *Aesculus hippocastanum* L. (horse chestnut) bark.

**Material and Method:** Bioactive compounds were extracted from *A. hippocastanum* bark using water and ethanol as solvent. The extracts were tested against 10 clinical strains isolated from urine of patients diagnosed positive to urinary tract infections and including five Gram-positive bacteria (*Kocuria rhizophila* 1542, *Enterococcus avium* 1669, *Staphylococcus simulans* 5882, *Conybacterium spp* 1638, *Enterococcus faecalis* 5960) and five Gram-negative (*Proteus mirabilis* 1543, *Morganella morganii* 543, *Citrobacter freundi 426*, *Acynetobacter baumannii* 5841 and *Achromobacter xylosoxidans* 4892). *Staphylococcus aureus* ATCC 6538 and *Escherichia coli* ATCC 25922 were used as standard Gram+ and Gram-respectively. The susceptibility of the test strains to antibiotic was assessed using the Kirby Bauer disc diffusion method while the antibacterial activity of the extracts was evaluated using the well diffusion method. We finally determined the Minimum inhibitory concentration (MIC) and the minimum bactericidal concentrations (MBC) by the microdilution method.

**Results:** *A. hippocastanum* bark possessed a dry matter content of 65.73%. The volume yield of the ethanolic and aqueous extract (AE) was 77,77% and 74,07% (v/v), respectively, whereas their mass yields were 13,4% and 24,3% (w/w), respectively. All the bacteria were susceptible to amoxiclav, imipenem and ceftriaxone, both standard bacteria (*E. coli* ATCC 25922 and S. aureus ATCC 6538) were sensitive to all antibiotics while the clinical strains were resistant to at least one antibiotic. *K. rizophilia* 1542 and *Conybacterium* spp 1638 were the most resistant bacteria both with multidrug resistance index of 0.45. Except AE on *P. Mirabilis* 1543 and *E. faecalis* 5960 (0 mm), both AE and EE were active against all the microorganisms tested with inhibition diameters (mm) which ranged from 5.5-10.0 for AE and 8.0-14.5 for EE. The MICs of EEs varied from 1-4 mg/ml while those of EAs varied from 4-16 mg/ml. Almost all the MBCs of AEs were indeterminate (>64 mg/ml) while those of EE were successfully determined. The ethanolic extracts (EE) were overall more active than the aqueous ones.

**Conclusion:** The *A. hippocastanum* bark extracts had overall weak antibacterial activity (MIC ≥0.625 mg/ml) and bacteriostatic potential (MBC/MIC ≥16) on both Gram-positive and Gram-negative bacteria. Therefore, studies with other solvents (such as methanol and chloroform), other extraction techniques, and synergy tests with conventional antibiotics are needed to conclude on a potential better antimicrobial activity of this plant material.

## Introduction

Urinary tract infections (UTIs) are very common infections and occur at least once in a lifetime. These infections are serious public health issues and are responsible for nearly 150 million disease cases every year worldwide [1]. UTIs are defined as any infection, commonly of bacterial origin, which occurs in any part of the urinary system [1]. When UTIs are localized in urethra, they are called urethritis, cystitis (when localized in the bladder), pyelonephritis (infection of the kidneys), and vaginitis (infection of the vagina) [2,3]. UTIs are more likely to occur in women, because, compared to men, their urethra is shorter and there is relative proximity between the urethra and the anus [4]. In addition, the prevalence of UTIs among sexually active young women has been reported to vary from 0.5 to 0.7 person per year, while this incidence rate among young men was only 0.01 [5]. Otherwhise, the so-called uropathogenic bacteria is responsible for 80-90% of UTIs but other germs such as *Staphylococcus saprophyticus, Pseudomonas aeruginosa, Staphylococcus aureus, Klebsiella pneumoniae, Proteus mirabilis, Acinetobacter baumannii, Streptococcus,* and *Enterococcus faecalis* are also sometimes involved [6]. This pathology is very often and easily managed with antibiotics [7] such as trimethoprim-sulfamethoxazole (TMP-SMX), nitrofurantoin, or Fosfomycin for 3–5 days [8] and sometimes by cephalosporins (fluoroquinolones; cefixime) and β-lactams (amoxicillin-clavulanate) [9]. However, unfortunately, resistance to antibiotics, which spares no area, has made UTIs more difficult to treat. Indeed, several recent studies have reported a considerable increase in antibiotic resistance in uropathogens and an unprecedented number of multidrug-resistant germs worldwide [10,11]. To respond to this dangerously worrying situation, research teams from all over the world are permanently evaluating the use of potential alternatives to antibiotics such as nanoparticles [12], probiotics [13,14], phage therapy [15], antimicrobial peptides [16] and medicinal plants [17,18]. Among these alternatives, medicinal plants appear as the most credible [19] since it is well known that plants have been used for millennia in the treatment and prevention of various diseases, including bacterial infections and some of these herbal remedies have been shown to be effective in preventing and treating UTIs [19]. In this context, certain plants such as *Aesculus hippocastanum* which are very widespread (and therefore available) but very few used deserve to be investigated for their antimicrobial properties.

Therefore, the aim of this work was to assess the antibacterial properties of ethanolic and aqueous *A. hippocastanum* bark extract against some bacteria involved in urinary tract infections.

## 2- Material and method

### 2-1- Vegetal material

The plant material used in this study was the bark of *Aesculus hippocastanum*. The plant was collected in June 2021 near the Medical Institute of the Peoples’ Friendship University of Russia (MG23+9H Obroutchevski, Moscow,Russia). After harvest, the plant was taken directly to the laboratory where it was dried at 37°C until the weight was constant. The dry matter content was calculated; the plant was grinded and the powders with particle sizes lower than 1 mm were stored in a sterile airtight container until further use.

### 2-2- Bacterial strains

The microorganisms used for the screening of antimicrobial activity consisted of 10 bacteria (five Gram positive bacteria and 5 Gram negative) isolated from urine of patients diagnosed positive to urinary tract infections. The five Gram + were *Kocuria rhizophila* 1542, *Enterococcus avium* 1669, *Staphylococcus simulans* 5882, *Conybacterium spp* 1638 and *Enterococcus faecalis* 5960 while the five Gram - were *Proteus mirabilis* 1543, *Morganella morganii* 543, *Citrobacter freundi 426, Acynetobacter baumannii* 5841 and *Achromobacter xylosoxidans* 4892. *Staphylococcus aureus* ATCC 6538 and *Escherichia coli* ATCC 25922 were used as standard Gram+ and Gram-respectively. All strains were provided by the Department of Microbiology and Virology of the Peoples’ Friendship University of Russia.

### 2-3- Chemicals and media

Dimethyl sulfoxide (DMSO) was purchased from BDH Laboratories, VWR International Ltd., USA. We also used BHIB (Brain Heart Infusion Broth) (HiMedia™ Laboratories Pvt. Ltd., India), Muller Hinton Agar (MHA HiMedia™ Laboratories Pvt. Ltd., India), Sabouraud Dextrose Broth (SDB, HiMedia™ Laboratories Pvt. Ltd., India) and all other reagents and chemicals used were of analytical grade.

### 2-4- Extraction of active compounds

Ethanolic solution (80%, v/v) and distilled water was used because they have been reported to be an efficient solvent for the extraction of bioactive compounds in the medicinal plants used in this study. As we described in our previous investigation [18] fifty grams (30g) of vegetal material was weighed and added to 270 ml of the solvent in separate conical flasks. The flasks were covered tightly and were shaken at 200 rpm for 24h and 25°C in a shaker incubator (Heidolph Inkubator 1000 coupled with Heidolph Unimax 1010, Germany). The mixtures were then filtered by vacuum filtration, using Whatman filter paper *№* 1 then concentrated at 40°C in rotary evaporator (IKA RV8) equipped with a water bath IKA HB10 (IKA Werke, Staufen, Germany) and a vacuum pumping unit IKA MVP10 (IKA Werke, Staufen, Germany). To avoid losses, the extracts were collected when the volumes were small enough and placed in petri dishes previously weighed and then incubated open at 40°C until complete evaporation. The final dried crude extracts were weighed. Extraction volume and mass yield were determined using the following formulas:

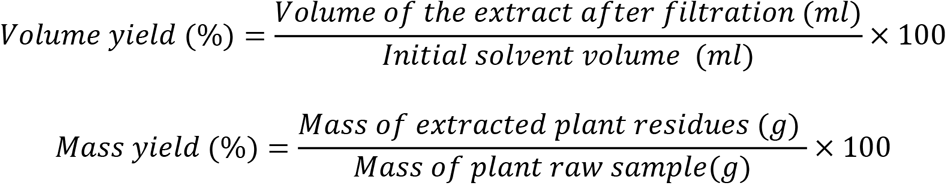

### 2-5- Preparation of antimicrobial solution

For each plant extract, the crude extract was dissolved in the required volume of DMSO (5%, v/v) to achieve a concentration of 128 mg/ml. The extracts were sterilized by microfiltration (0.22 μm) and the solution obtained was used to prepare the different concentrations used in the analytical process.

### 2-6- Inoculum preparation

Bacteria were cultured for 24 h at 37 °C in 10 ml of BHI broth. After incubation, the cells were collected by centrifugation (7000 g, 4 °C, 10 min), washed twice with sterile saline, resuspended in 5 mL of sterile saline to achieve a concentration equivalent to McFarland 0.5 using DEN-1 McFarland Densitometer (Grant-bio).

### 2-7- Evaluation of susceptibility of the test bacteria to antibiotics

The modified Kirby–Bauer’s disk method described in our previous study [20] was used to study the antibiotic sensitivity of the bacterial strains, and the following eight antibiotics disks were used: amoxicillin, 30 μg/disk; ampicillin, 25 μg/disk; cefazolin, 30 μg/disk; cefazolin/clavulanic acid, 30/10 μg/disk; 30 μg/disk; ceftriaxone, 30 μg/disk; ciprofloxacin, 30 μg/disk; Fosfomycin, 200 μg/disc; imipenem (IMP), 10 μg/disc; nitrofurantoin, 200 μg/disk; tetracyclin (TE), 30 μg/disc and trimethoprim, 30 μg/disk. The inhibition diameters were measured and interpreted referred to the Clinical & Laboratory Standards Institute [21]. Resistance R, Intermediate I, and Sensitive S interpretations were obtained automatically using algorithms written in Excel software [Microsoft Office 2016 MSO version 16.0.13628.20128(32 bits), USA] with the parameters described in Table 1 [10].

**Table 1:**
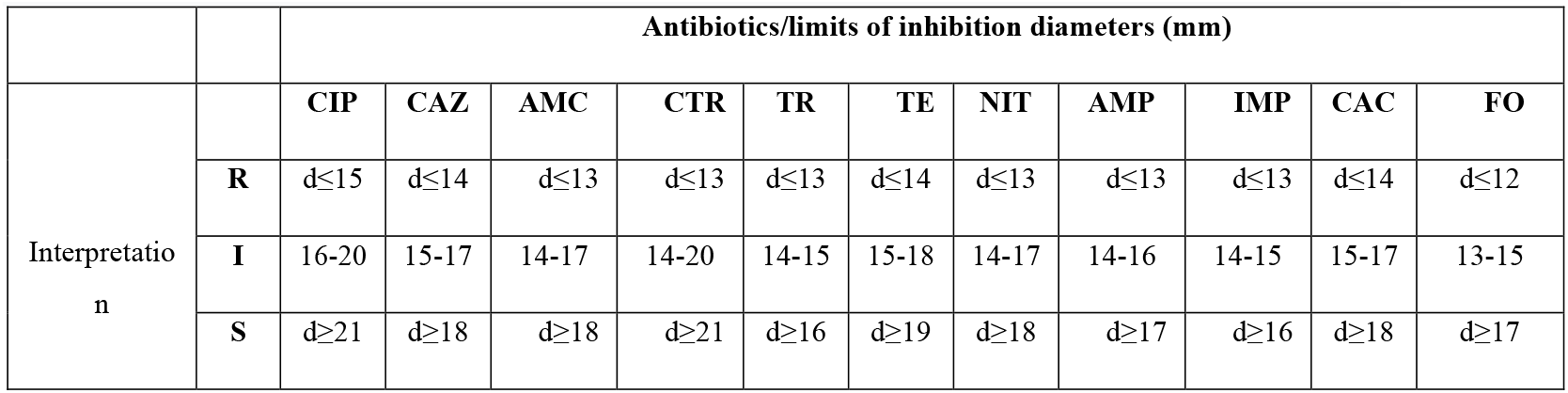
Interpretation criteria for antibiotic sensitivity[10, 21]; Amoxycillin (AMC), Ampi5cillin (AMP), Cefazolin (CZ); Cefazolin/ clavulanic acid (CAC); Ceftazidime (CAZ); Ceftriaxone (CTR); Ciprofloxacin (CIP); Fosfomycin (FO); Imipenem (IMP); Nitrofurantoin (NIT); Tetracyclin (TE) and Trimethoprim (TR).

|  |  | Antibiotics/limits of inhibition diameters (mm) |  |  |  |  |  |  |  |  |  |  |
| --- | --- | --- | --- | --- | --- | --- | --- | --- | --- | --- | --- | --- |
|  |  | CIP | CAZ | AMC | CTR | TR | TE | NIT | AMP | IMP | CAC | FO |
| Interpretation | **R** | d≤15 | d≤14 | d≤13 | d≤13 | d≤13 | d≤14 | d≤13 | d≤13 | d≤13 | d≤14 | d≤12 |
|  | **I** | 16-20 | 15-17 | 14-17 | 14-20 | 14-15 | 15-18 | 14-17 | 14-16 | 14-15 | 15-17 | 13-15 |
|  | **S** | d≥21 | d≥18 | d≥18 | d≥21 | d≥16 | d≥19 | d≥18 | d≥17 | d≥16 | d≥18 | d≥17 |

### 2-8- Screening of antibacterial activity

#### 2-8-1- Assessment of antimicrobial activity using well diffusion method

The well diffusion method previously described [18] was used to assess the antimicrobial activity of the extracts. Briefly, 15 ml of sterile Muller Hinton Agar (for bacteria) was poured into petri dishes and 100 μl of each microorganism were spread. Wells with a capacity of 20 μl were drilled on the culture medium and 20 μl (at 128 mg/ml) of each plant material was added. The sterile DMSO 5% used to prepare the extracts was used as negative control and all the trials was done in triplicate. After incubation at 37 °C for 24 h, the inhibition diameters were measured.

#### 2-8-2- Determination of minimum inhibitory concentrations (MIC)

MIC is the lowest concentration of antibacterial agent that completely inhibits the visible bacterial growth. The MIC of the extracts was determined using the microbroth dilution method as previously described without any modification [22]. Briefly, 100 μL of broth (BHIIB) was added to all the wells of sterile U-bottom 96-well microplates and extracts preparations (128mg/ml) were subjected to serial twofold dilution. Each column represented one type of extract and a single strain. DMSO 5% was used as negative control. For each test well, 10 μL of the respective inoculum was added (with turbidity equivalent to a 0.5 McFarland scale). Finally, the plates were covered and incubated at 37°C for 24 h and after incubation, MIC was considered the lowest concentration of the tested material that inhibited the visible growth of the bacteria.

#### 2-8-3- Determination of minimum bactericidal concentration (MBC)

MBCs were determined by subculturing the wells without visible growth (with concentrations ≥ MIC) on MHA plates. Inoculated agar plates were incubated at 37°C for 48h and MBC was considered the lowest concentration that did not yield any microbial growth on agar.

#### 2-8-4-Tolerance level

Tolerance level of tested bacterial strains against aqueous and ethanolic extract was determined using the following formula:

Tolerance = MBC/MIC

The characteristic of the antibacterial activity of extracts was determined by the tolerance level indicating the bactericidal or bacteriostatic action against the tested strains. When the ratio of MBC/MIC is ≥16, the antibacterial efficacy of the test agent is considered as bacteriostatic, whereas MBC/MIC ≤4 indicates bactericidal activity [23].

## 3- Results and discussion

### 3-1- Dry matter and extraction yield

The dry matter, water content and ethanolic (EE) and distilled water extract (AE) yields of *Aesculus hippocastanum* bark are presented in table 2. We found that *A. hippocastanum* bark possessed a dry matter content of 65.73%. Dry matter is an indicator of the amount of constituents (excluding water) in the sample tested. The drying allowed to remove water from the plant in order to standardize the extraction and to make this work reproducible if other authors from another region project to work with the same plant. Furthermore, we obtained an extraction volume yield of 74,07% with distilled water as solvent and 77,77% with ethanol. The difference in extraction volume yields can be explained by losses during the extraction process. Indeed, as reported by [18] we noticed that the filtration of ethanolic extracts was faster (less than 5 minutes for 300 ml) compared to the aqueous extract which took much longer (more than 50 minutes on average for 300 ml) and required in average 6 filter change. Despite the filtration, the EAs still looked cloudy while the EEs were completely clear. This residue cloudiness could therefore explain the higher mass yield in AE (24,3%) compared to EE (13,4%). Several authors [17,18, 24, 25] reported observations similar to our findings while others [26,27] found opposite results. Therefore, as Arsene et al. [18] pointed out in their previous work, the extraction performance depends on several factors, including the method use, extraction time, the solvents, and the equipment used. In addition, Mouafo et al. [17] reported that the high yields of phytoconstituent does not necessarily imply better antimicrobial activity and this was further assessed in the present work by evaluating the antimicrobial activity of each of the extracts using the well diffusion method and the determination of minimum inhibitory concentrations (MIC) and minimum bactericidal concentrations (MBC).

**Table 2:** Dry matter, water content, volume and mass extraction yields of *A. hippocastanum bark*

|  | Aqueous extract | Ethanolic extract |
| --- | --- | --- |
| Dry matter (%) | 65,73 |  |
| Water content (%) | 34,27 |  |
| Volume yield (%) | 74,07 | 77,77 |
| Mass yield (%) | 24,3 | 13,4 |

### 3-2- Susceptiility of clinical strains to antibiotics

The susceptibility of the tested bacteria to eleven (11) antibiotics was evaluated by the Kirby Bauer disc method, the Multidrug resistance Index (MDR) was calculated, and the results were reported in Table 3. As shown in the table 3, no bacteria were resistant to amoxiclav, imipenem and ceftriaxone while 7/12 were resistant to ampicillin, 6/12 to trimethoprim, 5/12 to tetracycline and ceftazidime/clavulanic acid, 4/12 to ceftazidime, 3/12 to fosfomycin, 2/12 to nitrofurantoin and 1/12 to ciprofloxacin. Interestingly, both standard bacteria (*E. coli* ATCC 25922 and *S. aureus* ATCC 6538) were sensitive to all antibiotics while the clinical strains were resistant to at least one antibiotic. Among clinical strains, MDRs ranged from 0.09 to 0.45. *K. rizophilia* 1542 and *Conybacterium* spp 1638 were the most resistant bacteria with MDRs of 0.45 each. The results observed with these clinical strains are similar to those reported in other studies [10,11]. While the strongest resistance was observed on ampicillin (which is an antibiotic from the beta-lactam family, from the group A of the penicillins), no resistance was observed in amoxiclav which is a derivative of amoxicillin (antibiotic of the beta-lactam family, of the aminopenicillin group) of the same family, with the only difference that the latter is combined with clavulanic acid (beta-lactamase inhibitor). This shows that antibiotic adjuvants can also play an important role in combating resistant germs [28]. Furthermore, similarly to our findings, other studies have reported low resistance of uropathogens to imipenem and ceftriaxone [11,29]. In addition, the sensitivity of all strains to amoxiclav, imipenem and ceftriaxone demonstrates that these 3 antibiotics are effective on a wide range of bacteria, including clinical strains with high resistance to other antibiotics. Therefore, these antibiotics which have kept a good activity against various microorganisms should be carefully used and administered only under prescription and after a prior antibiogram.

**Table 3:** Susceptibility to antibiotics of the tested bacteria. Amoxycillin (AMC), Ampi5cillin (AMP), Cefazolin (CZ); Cefazolin/ clavulanic acid (CAC); Ceftazidime (CAZ); Ceftriaxone (CTR); Ciprofloxacin (CIP); Fosfomycin (FO); Imipenem (IMP); Nitrofurantoin (NIT); Tetracycline (TE) and Trimethoprim (TR). R= Resistant, I= Intermediate and S=Sensitive

|  |  | NIT | TE | CTR | AMC | FO | CAZ | IPM | CAC | CIP | AMP | TR | MDR |
| --- | --- | --- | --- | --- | --- | --- | --- | --- | --- | --- | --- | --- | --- |
| Gram + | *K. rizophila* 1542 | 10±0(R) | 13±0(R) | 22±0(S) | 22±1(S) | 28±2(S) | 10±0(R) | 23±1(S) | 6±0(R) | 30±1(S) | 13±1(R) | 21±2(S) | 0,45 |
|  | *E. avium* 1669 | 21±1(S) | 6±0 (R) | 30±4(S) | 25±3(S) | 31±3(S) | 23±1(S) | 27±4(S) | 24±2(S) | 15±0(R) | 6±0(R) | 6±0(R) | 0,36 |
|  | *S. simulans* 5882 | 23±1(S) | 20±2(S) | 19±1(S) | 27±3(S) | 38±4(S) | 11±0(R) | 34±3(S) | 11±0(R) | 26±2(S) | 6±0(R) | 20±2(S) | 0,27 |
|  | *Conyobacterium spp* 1638 | 20±0(S) | 32±4(S) | 32±5(S) | 36±2(S) | 14±1(R) | 6±0(R) | 27±2(S) | 6±0(R) | 26±1(S) | 9±0(R) | 10±0(R) | 0,45 |
|  | *E. faecalis* 5960 | 21±1(S) | 11±0(R) | 22±1(S) | 27±1(S) | 31±3(S) | 17±1(I) | 24±2(S) | 6±0(R) | 32±3(S) | 6±0(R) | 6±0(R) | 0,36 |
|  | *S. aureus* ATCC 6538 | 22±1(S) | 32±3(S) | 30±2(S) | 35±4(S) | 38±2(S) | 25±1(S) | 28±3(S) | 25±2(S) | 32±0(S) | 20±1(S) | 31±2(S) | 0,00 |
| Gram - | *P. mirabilis* 1543 | 19±1(S) | 6±0(R) | 30±4(S) | 25±2(S) | 31±0(S) | 23±2(S) | 27±1(S) | 24±1(S) | 30±2(S) | 16±1(I) | 6±0(R) | 0,18 |
|  | *M. morgani* 1543 | 15±0 (I) | 6±0(R) | 33±2(S) | 17±1(I) | 13±0(R) | 23±1(S) | 22±0(S) | 23±1(S) | 22±2(S) | 10±0(R) | 6±0(R) | 0,36 |
|  | *C. freundi* 426 | 21±1 (S) | 30±3(S) | 27±2(S) | 35±1(S) | 40±2(S) | 12±0(R) | 37±1(S) | 10±0(R) | 30±2(S) | 6±0(R) | 22±2(S) | 0,27 |
|  | *A. baumannii* 5841 | 26±2(S) | 16±0(I) | 27±3(S) | 35±1(S) | 32±3(S) | 19±2(S) | 34±2(S) | 17±1(I) | 30±1(S) | 21±2(S) | 14±0(R) | 0,09 |
|  | *A. xylosoxidans* 4892 | 6±0(R) | 11±0(R) | 23±2(S) | 36±4(S) | 6±0(R) | 16±0(I) | 32±3(S) | 16±1(I) | 20±2(I) | 20±1(S) | 6±0(R) | 0,36 |
|  | *E. coli* ATCC 25922 | 21±3(S) | 30±2(S) | 24±1(S) | 20±1(S) | 40±4(S) | 18±1(I) | 32±2(S) | 16±1(I) | 24±0(S) | 26±0(S) | 24±0(S) | 0,00 |

### 3-4- Inhibition zone of the extracts against the tested bacteria

Figure 1 presents the inhibition diameters of ethanolic (EE) and aqueous (AE) extract of *Aesculus hippocastanum* bark on the tested microorganisms. Except AE on *Proteus Mirabilis* 1543 and *Enterococcus faecalis* 5960 (0 mm), both AE and EE were active on all microorganisms tested with inhibition diameters (mm) which ranged from 5.5-10.0 for AE and 8, 0-14.5 for EE. The ethanolic extracts (EE) were overall more active than the aqueous ones. Consequently, this means that ethanol extracts more compounds with antimicrobial properties compared to water although we found above that the extraction yields with water were higher than that with ethanol. Several authors have reported that compounds with antimicrobial activity such as flavonoids, polyphenols, tannins and alkaloids are generally insoluble in water but soluble in ethanol[30,31]. Other authors such as Arsene et al.[18], Mouafo et al. [17] and Evbuomwan et al. [32] also pointed out that ethanol extracts more antimicrobial compounds from plant materials opposed to water. In addition, we found that extracts of *A. hippocastanum* bark were both active against Gram + bacteria as against Gram - bacteria. To our knowledge, the antibacterial properties of this plant have not yet been investigated but our findings suggest that *A. hippocastanum* bark has constituents exhibiting a broad-spectrum antimicrobial.

**Figure 1:**
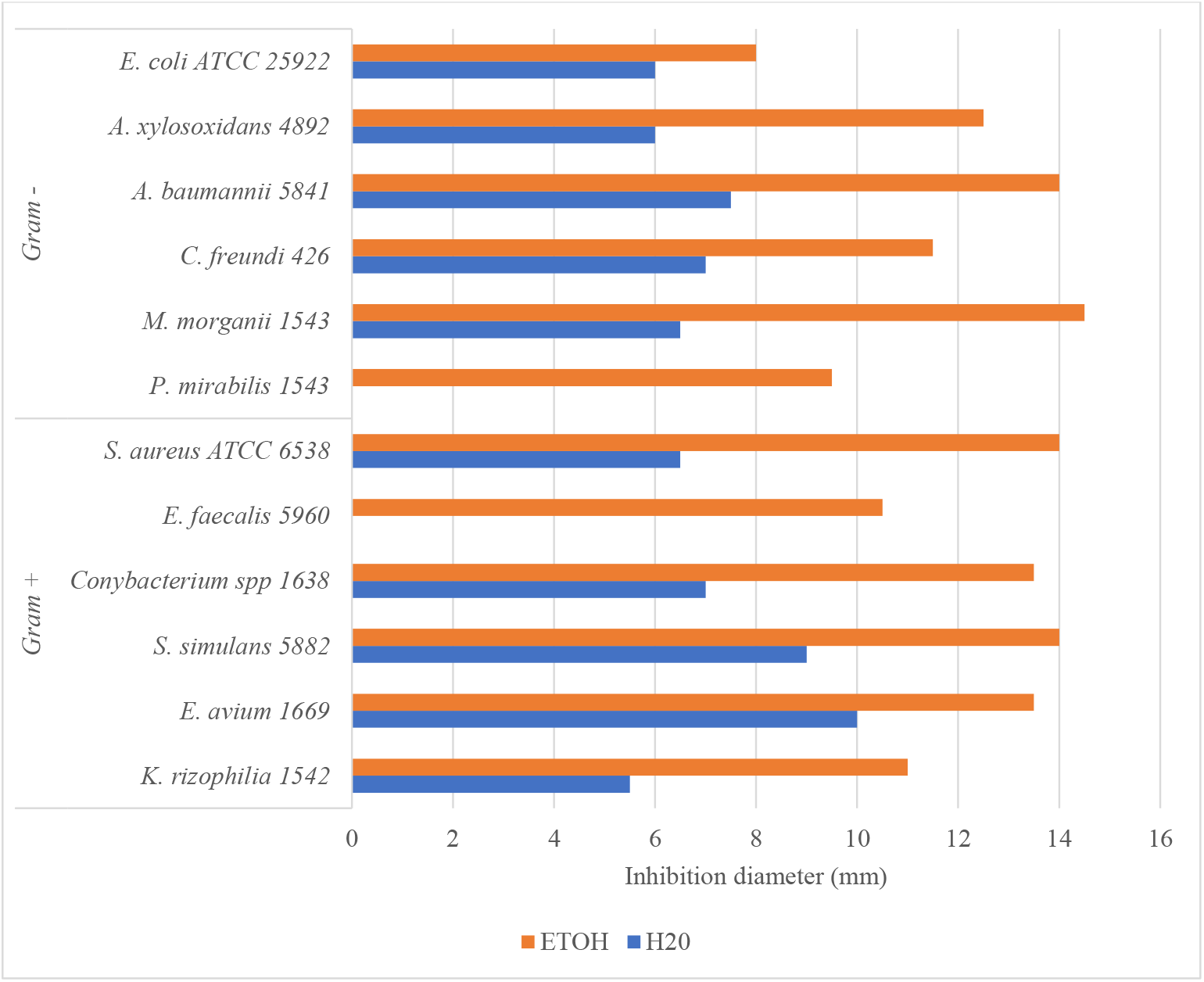
Inhibition diameter of ethanolic and aqueous Aesculus hippocastanum bark extract on the tested bacteria

Most of the research conducted on *A. hippocastanum* focused on its properties to regulate circulatory system imbalances and relieve attacks of hemorrhoids [33]. In a study conducted by Owczarek et al. [33] on the composition of *A. hippocastanum* bark, it has been reported that this plant presents mainly two groups of compounds: coumarins and proanthocyanidins. Among the coumarins there was mainly esculin (over 17%) and fraxin (over 7%) while among the proanthocyanidins, the authors found epicatechin (over 6%) and procyanidin A2 (over 5%) (Figure 2) [33]. Owczarek et al. [33] finally conclude that, in total, about 40% of the *A. hippocastanum* bark extract could be attributed to simple phenolics compounds detectable by LC-PDA. Unfortunately, we were unable to assess the composition of our extracts in order to compare with the data in the literature. However, we can hypothesize that the antimicrobial properties of *A. hippocastanum* bark can be attributed to all its components or specifically to esculin [34], to fraxin[35] or proanthocyanidins[36] because these compounds (from other plants) have been reported to have antimicrobial properties. Notwithstanding our findings, further investigations are required to confirm or refute our hypothesis.

**Figure 2:**
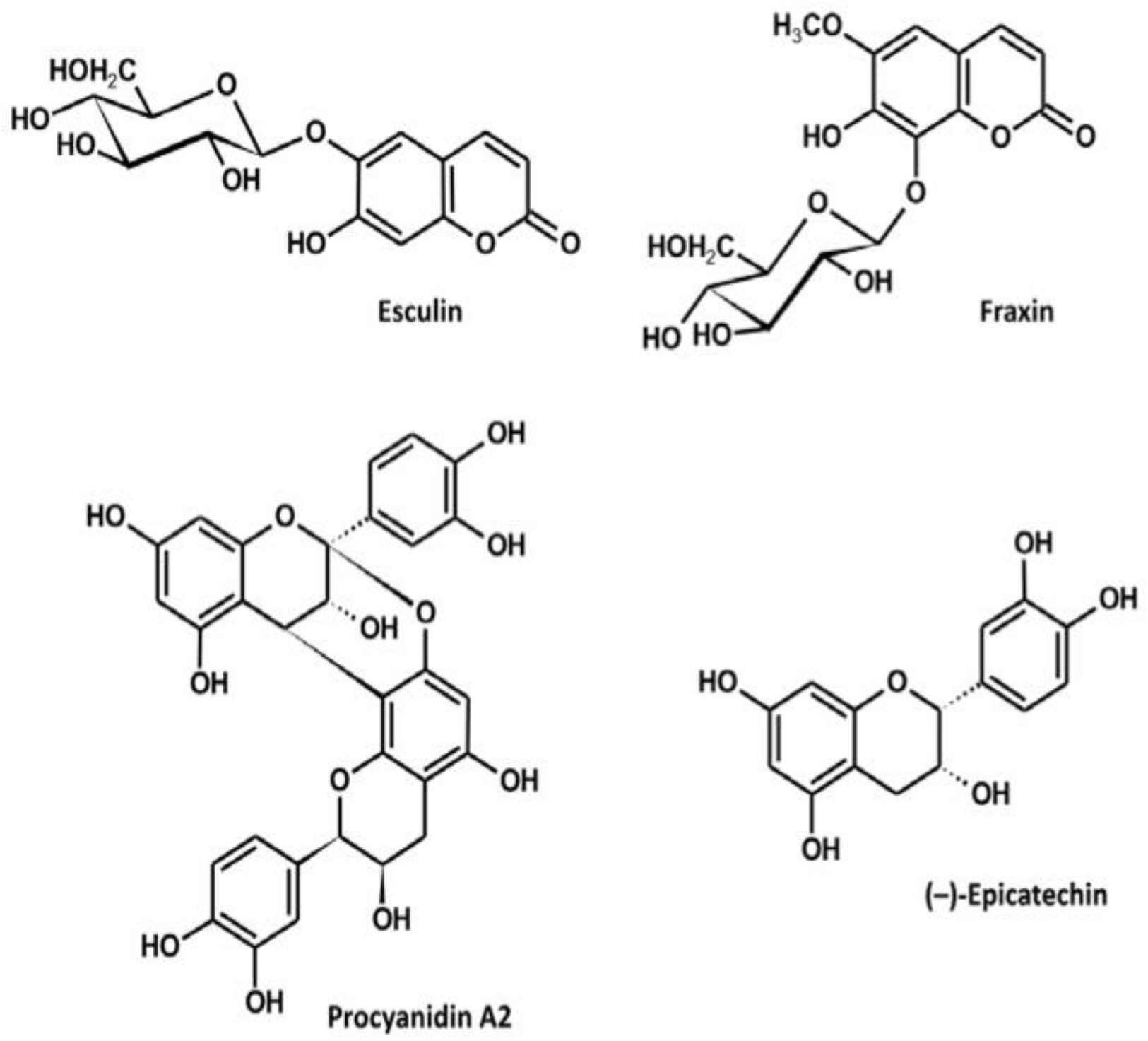
Structures of the major constituents of the A. hippocastanum bark[33].

### 3-4- Minimum inhibitory concentrations (MIC) and minimum bactericidal concentrations

After evaluating the antibacterial properties of aqueous (AE) and ethanolic (EE) extracts of Aesculus hippocastanum bark against the uropathogens tested using the well diffusion, we investigated the minimal inhibitory concentrations (MIC) and minimal bactericidal concentrations (MBC) of the two extracts. MIC and MBC are two very important elements in the search for new antimicrobials and respectively provide the minimum concentrations required to inhibit or kill the microorganism tested. Table 4 presents the MIC, the MBC, and the MBC/MIC ratio of our 2 extracts on the uropathogens investigated. Similarly, to the inhibition diameters, ethanolic extracts (EE) showed the best antimicrobial activity with the lowest MIC and MBC. The MICs of EEs varied from 1-4 mg/ml while those of EAs varied from 4-16 mg/ml. Almost all the MBCs of AEs were indeterminate (>64 mg/ml) while those of EE were successfully determined. With water as solvent, the highest antimicrobial activity was observed against *E. faecalis* 5960, *S. aureus* ATCC 6538 and *C. freundi* 426. Although the MIC and MBC of AE were high against these bacteria, AE was found to be bactericidal against *S. aureus* ATCC 6538 and *C. freundi* 426 since the MBC/MIC ratio was 4 for both bacteria. Indeed, Mondal et al. [23] reported that when the ratio MBC/ MIC is ≥16, the antibacterial efficacy of the test agent is considered as bacteriostatic, whereas MBC/MIC ≤4 indicates bactericidal activity. Similarly, although the MICs of EEs were relatively low, the MBCs of this extract were quite high, and we concluded that the antibacterial activity was overall bacteriostatic (MBC/ MIC ≥16) against most bacteria except against some Gram-positive bacteria such as *S. simulans* 5882, *Conybacterium* spp 1638 and *S. aureus* ATCC 6538. This difference in the activity (although not significant) of EE between Gram positive and Gram negative can be ascribed to the cell wall structure of Gram - bacteria, which differs from the structure of Gram + bacteria with a thin layer of peptidoglycan, the presence of a periplasm, and ease of exchange on the plasma membrane [18]. Furthermore, according to the classification of Kuete [37]), the different extracts could be considered as deserving a weak antimicrobial activity independently of the extraction solvent and the tested strain as they scored MIC value higher than 0.625 mg/ml. Therefore, further studies are needed to evaluate the antibacterial activity of *A. hippocastanum* bark extracts with other solvents and to assess their synergy with other antibiotics, as the weak antimicrobial activity observed here does not allow us to recommend our extracts plant as such for a possible use in the fight against antibiotic resistance and the management of urinary tract infections.

**Table 4:** Minimum inhibitory concentrations (MIC) and minimum bactericidal concentrations (MBC) of ethanolic and aqueous *Aesculus hippocastanum* bark extract on the tested bacteria

|  |  | Aqueous extract |  |  | Ethanolic extract |  |  |
| --- | --- | --- | --- | --- | --- | --- | --- |
|  |  | MIC<br>(mg/ml) | MBC<br>(mg/ml) | MBC/MIC | MIC<br>(mg/ml) | MBC<br>(mg/ml) | MBC/MIC |
| Gram + | *K. rizophilia* 1542 | 16 | >64 | - | 4 | >64 | - |
|  | *E. avium* 1669 | 4 | >64 | - | 2 | 64 | 32 |
|  | *S. simulans* 5882 | 16 | >64 | - | 2 | 16 | 8 |
|  | *Corynebacterium spp 1638* | 64 | 64 | 1 | 4 | 32 | 8 |
|  | *E. faecalis 5960* | 4 | 64 | 16 | 2 | 64 | 32 |
|  | *S. aureus ATCC 6538* | 16 | 64 | 4 | 2 | 16 | 8 |
| Gram - | *P. mirabilis 1543* | 8 | >64 | - | 2 | 64 | 32 |
|  | *M. morganii 1543* | 2 | >64 | - | 1 | 64 | 64 |
|  | *C. freundii 426* | 16 | 64 | 4 | 2 | 32 | 16 |
|  | *A. baumannii 5841* | 4 | >64 | - | 2 | 64 | 32 |
|  | *A. xylosoxidans 4892* | 4 | >64 | - | 2 | 64 | 32 |
|  | *E. coli ATCC 25922* | 16 | >64 | - | 2 | 16 | 8 |

## 4- Conclusion

The search for new antimicrobials is essential for overcoming antibiotic resistance in the management of bacterial diseases including urinary tract infections (UTIs). In this study we evaluated the antibacterial properties of aqueous and ethanolic extracts of *Aesculus hippocastanum* bark against ten (10) clinical uropathogenic bacteria and two (2) standard bacteria. The results showed that, except against few bacteria, the extracts had overall weak antibacterial activity (MIC≥0.625 mg/ml) and bacteriostatic potential (MBC/MIC ≥16) on both Gram positive and Gram-negative bacteria. Therefore, studies with other solvents (such as methanol and chloroform), other extraction techniques, and synergy tests with conventional antibiotics are needed to conclude on a potential better antimicrobial activity of this plant.

## Acknowledgement

This study has been supported by the RUDN University strategic Academic Leadership Program.

